# Fragment-Based Ligand Generation Guided By Geometric Deep Learning On Protein-Ligand Structure

**DOI:** 10.1101/2022.03.17.484653

**Authors:** Alexander S. Powers, Helen H. Yu, Patricia Suriana, Ron O. Dror

## Abstract

Computationally-aided design of novel molecules has the potential to accelerate drug discovery. Several recent generative models aimed to create new molecules for specific protein targets. However, a rate limiting step in drug development is molecule optimization, which can take several years due to the challenge of optimizing multiple molecular properties at once. We developed a method to solve a specific molecular optimization problem *in silico*: expanding a small, fragment-like starting molecule bound to a protein pocket into a larger molecule that matches that physiochemical properties of known drugs. Using data-efficient E(3) equivariant based neural networks and a 3D atomic point cloud representation, our model learns how to attach new molecular fragments to a growing structure by recognizing realistic intermediates generated *en route* to a final ligand. This approach always generates chemically valid molecules and incorporates all relevant 3D spatial information from the protein pocket. This framework produces promising molecules as assessed by multiple properties that address binding affinity, ease of synthesis, and solubility. Overall, we demonstrate the feasibility of 3D molecular structure expansion conditioned on protein pockets while maintaining desirable drug-like physiochemical properties and developed a tool that could accelerate the work of medicinal chemists.

## 1 Introduction

Drug discovery and design aims to find novel molecules to treat human disease. These molecules bind and control the activity of target proteins and other biomolecules in the human body. However, the drug discovery process is long and expensive. The vast space of possible molecules is repeatedly explored and filtered down into candidates most likely to be viable therapeutics. Moving the chemical search process from the benchtop to a laptop could significantly decrease the money, time, and human effort needed to create a new drug. Although many computational tools focus on the first step of identifying initial candidates, optimization of these molecules is a rate limiting step for drug discovery that can take years.

In this work, we focus on a specific molecular optimization task: expanding a small, fragment-like starting molecule bound to a protein pocket into a larger, more drug-like molecule. This task is a common challenge in molecule optimization (Schneider & Fechner, 2005). Fragment-based screening strategies (NMR, X-ray crystallography, virtual screening) and natural molecules (hormones, peptides) can be used to suggest starting orientations of fragments that bind to a 3D protein structure. Expanding these fragment-like molecules results in more desirable drug properties, such as higher affinity and specificity for the protein target from increased interactions with the protein binding site (Kuntz et al., 1999; Kenny, 2019; Hopkins et al., 2006). However, this task is challenging and typically costly for several reasons. Many properties must be optimized simultaneously some of which are themselves challenging to predict for both chemists and algorithms. These properties include binding to the target protein (affinity), solubility in water and fat, molecular size, and ease of synthesis. Molecules become “drug-like” when they fit into acceptable ranges of these properties (Bickerton et al., 2012). Additionally, chemical space is vast and unfeasible to enumerate fully; a search strategy is required to produce lists of useful candidates for medicinal chemists to utilize. AI agents can potentially learn to address each of these problems efficiently and take advantage of rapidly growing molecular datasets — unlike hand-crafted methods.

Unfortunately, current machine learning methods for molecule generation are ill-suited to this task. Many generative models aim to produce brand new molecules rather than explicitly expanding on an initial candidate ligand (ligands are molecules that can bind to proteins). Approaches based on variational autoencoders can produce diverse molecules related to input molecules, but struggled to produce molecules larger than the input (Masuda et al., 2020). Another challenge is that many methods do not generate ligands in the context of the 3D protein pocket and are evaluated based on molecule-only metrics, ignoring interactions with the protein. Recent breakthrough improvements in experimental techniques and computational tools like AlphaFold are making protein structures more readily available; this structural information should be utilized (Varadi et al., 2021).

To solve these problems, we present a framework for 3D, pocket-aware, ligand expansion inspired by behavioural cloning. First, we represent the generation process as a sequence of steps in 3D space, in which we attach new molecular fragments to the growing seed molecule. Our curated fragment library simplifies the possible actions while maintaining expressivity. In order to choose which fragment to add and the fragment geometry, we cast this sequence of steps as a supervised learning problem over state-action pairs. We create synthetic trajectories of state-action pairs by fragmenting ligands from a curated dataset of ligand-protein structures (“expert” molecules). We then learn to predict and score actions from states represented as 3D atomic point clouds using E(3) equivariant neural networks. This architecture allows us to incorporate the geometry of both the ligand and protein pocket. This robust approach requires no reward function, generates only valid molecules (following chemical rules such as proper valence), and incorporates all relevant 3D spatial information without separate encoding steps.

We evaluate the expanded ligands in terms of 12 molecular properties — several unique to our study — that address affinity, ease of synthesis, and hydrophobicity. We compare our performance to agents that choose actions based on state-of-the-art docking functions. Remarkably, our agent is able to generate ligands that match the property distributions of the expert molecules (the known ligands from our curated dataset). This is surprising given that our agents are never made aware of these final properties and are only trained on intermediate states. Our approach has several additional strengths. We are able to learn a complex task from a relatively small number of independent training examples (4000 protein-ligand pairs). Second, our approach is interpretable; the agent’s actions often align with a chemist’s intuition and basic physics. Thus, our learning framework and associated datasets may be useful for diverse tasks in ligand optimization.

## 2 Related Work

Recent work on molecular generation tasks highlight many of the challenges that these models face and the diversity of approaches. We discuss related work in the context of three categories: approaches to molecular representations and action spaces, incorporation of protein target information, and training objectives.

Representations and actions spaces have evolved over time in concert with advances in model architectures. In the earliest methods, usually based on hand-craft objectives, molecules were constructed in 3D space by attaching fragments, groups of atoms that make up common motifs in drug molecules (Schneider & Fechner, 2005). At that time, 3D structural data was more limited and ML architectures were less suited for these representations. The success of language models in other areas of machine learning led to use of string-based molecular representations, SMILES, that were constructed using RNNs (Bombarelli et al., 2018; Krishnan et al., 2021). Graphs are a more natural 2D representation for molecules, and many methods modeled the generation process as a sequence of adding nodes (atoms) or edges (bonds) using graph-neural networks (Li et al., 2018; De Cao & Kipf, 2018; Liu et al., 2018)). While SMILES and graph based representations benefit from large databases of molecules for training (CHEMBL, ZINC), it is harder to incorporate relative spatial information about the protein target, which is 3D in nature. To leverage 3D information, several studies utilized atomic density grids; with this representation and CNNs, the entire ligand can be generated in “one step” by outputting a density over the voxelized 3D space (Masuda et al., 2020; Skalic et al., 2019). However, this approach requires a secondary optimization algorithm to fit molecules to the resulting density. Coming full circle, our approach harkens back to the original 3D fragment concept but leverages recent advances in architectures that act on 3D point clouds. DeepFrag (Green et al., 2021) similarly expands a parent ligand by adding a fragment; but whereas Deepfrag predicts only a single 2D fragment to add to a pre-selected location, we attempt a more complex sequence of additive actions in 3D with no restriction on location.

The 3D structure of the target protein is another potential source of information for ligand generation and optimization tasks. Many molecular generation methods do not use protein target information, aiming only to sample chemical space. Some methods, such as Seq2Mol, rely only on the 1D sequence of amino acids which is incorporated as a sequence embedding (Ghanbarpour & Lill, 2020). To take advantage of the 3D shape of the binding pocket, methods have encoded the protein atom or residue graph into a latent vector, which is then used to condition the generation process (Drotár et al., 2021; Krishnan et al., 2021). Masuda et al. (2020) utilize 3D density grids and a 3D CNN to encode the protein pocket into a latent space, though the weights are separate from the ligand encoder. Our approach captures both the ligand and protein atoms simultaneously in the point cloud, distinguishing them with one-hot encoded features, which may better capture relative geometry.

Finally, approaches differ in whether they aim to reconstruct example molecules, to optimize particular properties, or both. Reinforcement learning approaches attempt to generate molecules with a particular optimized property, such as high affinity. MoleGuLAR (De Cao & Kipf, 2018) use multiple metrics and alternates the reward during the learning process for better stability. However rewards must be considered with caution, as algorithms to predict molecular properties such as binding affinity are themselves imperfect. In contrast, reconstruction focused methods, usually conditional VAEs or GANs, aim to imitate molecules in a database, without explicitly incorporating metrics or properties. Our approach is similar, but trains an agent to imitate individual actions in the generation process, which may be easier to learn and generalize than an objective based only on reproducing the complete ligand.

## 3 Methods

### 3.1 Action Space

The core of our framework is an agent which selects actions to take based on the current molecular state. The initial state is the 3D atomic structure (atom xyz locations, elements, bonds) of the seed ligand molecule and the protein pocket. The agent sequentially adds fragments to the ligand, connected by single bonds, until the molecular weight of the ligand hits a user-specified goal (Figure 1). These fragments are selected from a library of 28 common functional groups found in ligands (Figure 9). Each action is broken down into two steps (1) selecting a location to attach a fragment (2) given a location, choosing which fragment to add and the dihedral angle specifying the attachment geometry. Two separate models are trained to make predictions for each step.

**Figure 1:**
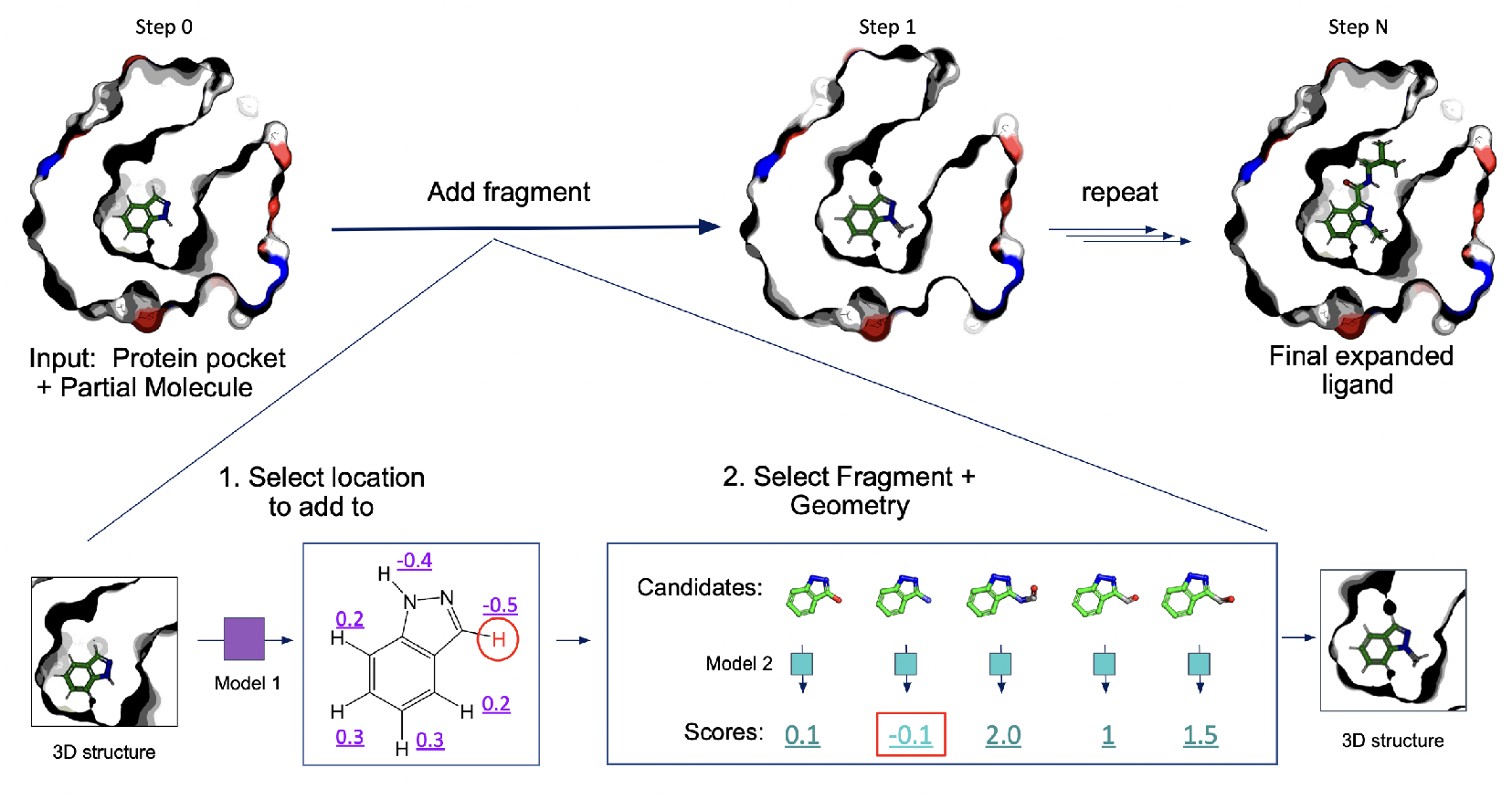
Ligand generation through sequential additional of fragments guided by geometric deep learning models. Generation is instantiated with a seed fragment placed inside of the protein pocket of interest. The method sequentially adds fragments to the ligand, connected by single bonds, until the molecular weight of the ligand hits a user-specified goal. Each action is broken down into two steps. First, to select a location to attach a fragment, potential attachment points are assigned a score by model 1 (purple). After selecting the best scoring location (red circle), we generate a set of candidate structures by sampling fragments and dihedral angles specifying the attachment geometry. These candidates are scored by model 2 (cyan). The best scoring state is selected (red box) and the process is repeated.

For the location step, ligand hydrogen atoms correspond to the possible locations at which new fragments can be attached (the hydrogen atom will be replaced by the fragment). Thus, the model is trained as a binary classifier to output scores for each ligand hydrogen atom corresponding to whether the atom should be an attachment point or not. This model takes as input both the current ligand state and protein pocket. For the fragment step, the model is trained to output a single scalar score given a ligand with candidate attached fragment and the protein pocket. The candidate fragment atoms to evaluate are flagged using the atom feature vectors. All candidates (across fragments and dihedral angles) are created and we use the model scores as a heuristic to rank these candidate states.

This process can be run to output a single ligand, by greedily choosing the highest scoring actions at each state. Alternatively, we can use the model scores and ranked actions as heuristics to inform more sophisticated search strategies that output a set of diverse ligands. In this work, we focus on the greedy case.

### 3.2 Architecture

To predict actions from atomic structures, we use E(3) equivariant neural networks, which act on 3D points clouds (Thomas et al., 2018; Geiger et al., 2020). Previous works by Eismann et al. (2020a;b); Townshend et al. (2021) have shown success for end-to-end learning from 3D atomic structures. This point representation in 3D space allows us to represent the relative positioning of the atoms in the protein-ligand complex precisely, which is important to capture interaction between protein-ligand atoms. In addition, the inherent rotational-equivariance property of these networks allows more efficient learning with the relatively small datasets of protein-ligand complex structures.

Our architecture has two main components: (1) embedding unit and (2) aggregator unit (Figure 2). The embedding unit consists of two layers of sequential application of self-interaction, point-convolution, self-interaction (Schütt et al., 2017), nonlinearity, and point normalization (Eismann et al., 2020b), which aims to learn embeddings of the input structure. Each atom/point in 3D is associated with a feature vector. At input, they are the basic element type of the atom (C, O, N, P, S, H and F/Cl/Br) encoded as a one-hot vector and Boolean flags indicating whether an atom belongs to the ligand, protein, or the attached fragment when applicable. The point-wise feature vectors are updated through the embedding unit layers by aggregating local information of the nearest *N* neighboring points.

**Figure 2:**
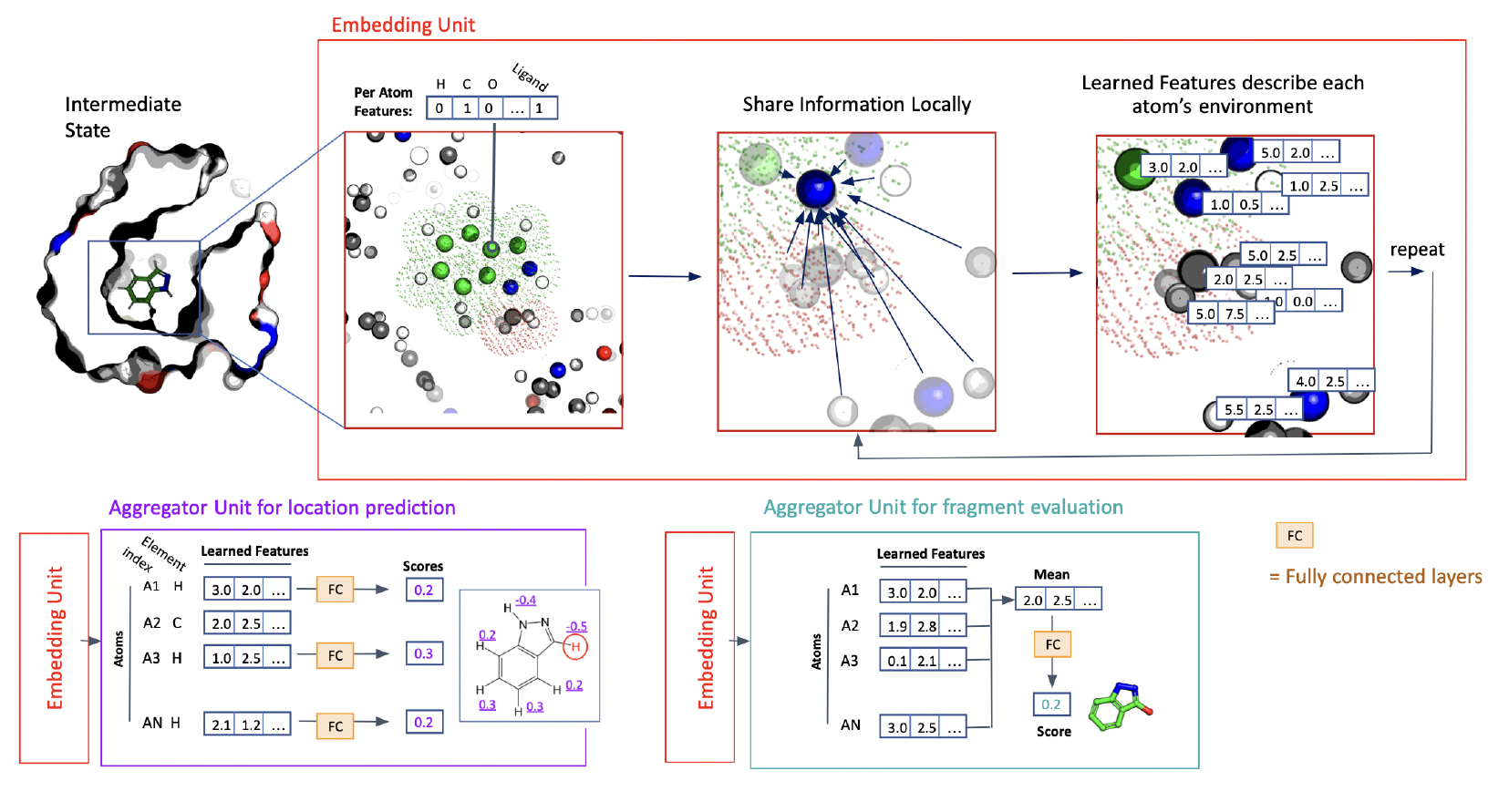
Model architecture. Each model consists of an embedding unit and aggregator unit. The embedding unit captures information about the chemical environment; each atom’s feature vector is updated based on the features vectors of the closest *N* atoms. The aggregator unit is specific for each task, whether the location prediction task or the fragment evaluation task. FC consists of a sequence of 1 fully connected layer followed ELU activation and 1 final fully connected layer. Ligands from existing protein-ligand complexes were used to train these networks.

The learned embedding vectors of the final layer of the embedding unit act as input to the aggregator unit. The aggregator unit consists of 2 fully connected (FC) layers followed by ELU activation function except for the final FC layer (Clevert et al., 2016). The aggregation process differs depending on the type of actions being evaluated. To score a candidate fragment at a given location, we take the average of the learned feature vectors over all points to give one mean feature vector, which is then passed to the aggregator unit. The output of the aggregator unit is a scalar value in this case.

The other type of action is choosing a location to add a fragment. For this case, we instead select only the atoms corresponding to hydrogen atoms of the ligand, concatenate the corresponding feature vectors, and pass it to the aggregator unit. The final output is a vector of scores corresponding to each ligand hydrogen atom.

For details on the hyperparameters used for each component in the architecture, please refer to Supplement A.

### 3.3 Datasets

Ligands were derived from the PDBBind 2019 refined dataset (Liu et al., 2015), a collection of proteinligand structures with high resolution. The dataset was filtered to remove common biomolecules (lipids, peptides, carbohydrates, and nucleotides) and any compounds outside a molecular weight range to restrict the molecules to those that have similar physiochemical properties to known drugs. Synthetic “expert” sequences of states (*s*_0_…*s*_*i*_…*s*_*N*_) were created by sequentially removing fragments from each ligand using a custom graph matching algorithm until a minimum size was reach (this is then *s*_0_). Each removed fragment is replaced by a hydrogen atom in the intermediate states to satisfy valence requirements.

We then derived two datasets: a dataset used to evaluate the location to add fragments OPENBOND dataset), and a dataset to evaluate a specific candidate fragment (FRAGMENT dataset). We split both datasets using the same split of protein-ligand pairs into training (70%), validation (15%), and test (15%) sets. To prevent data leakage during training, we split these structures such that no protein in the test set had more than 30% sequence identity with any protein in the training and validation set. Benchmarking on expanded ligands (see 4.2) was performed on the test set which was not presented during training or validation.

For the OPENBOND dataset, we computed binary labels for each ligand hydrogen atom in the intermediate states (*s*_*i*_), corresponding to whether this hydrogen would be replaced by a fragment in the final state (*s*_*N*_). Because some of the sequences of states are much longer than others, we randomly selected a maximum of 6 labeled states for each ligand-protein pair to include in the dataset. For the FRAGMENT dataset, we selected up to 6 intermediate state (*s*_*i*_) and for each generated 1 near-native state that closely matched *s*_*i*+1_ (by root-mean square deviation of coordinates) and 6 decoy states with randomly sampled incorrect fragments or geometries. The near-native example is given a label of 1 and the decoy examples a label of 0. This resulted in significant augmentation of the dataset, with a total of over 100,000 examples (Table 1).

**Table 1:** Number of samples for each dataset presented during training.

| Task | Number of Samples |  |
| --- | --- | --- |
|  | Train | Val |
| OPENBOND | 1676 | 1030 |
| FRAGMENT | 105182 | 13885 |

### 3.4 Training and Evaluation

For both types of models described in 3.2, we formulate the training as a binary classification task. The binary cross entropy with logits loss between the actual and the predicted label is used as the loss function during training. To address issues with imbalanced datasets, as we have considerably more negative over positive samples, we randomly oversample the less frequent class respectively during training.

We train with the Adam optimizer in Pytorch (Paszke et al. (2019)) with learning rate of 0.01 and batch size of 8 for 30 epochs and monitor the loss on the validation set at every epoch. The weights of the best-performing network are then used to evaluate the predictions on the validation set. Please refer to Table 2 for performance on the validation set. We train the models on 1 NVIDIA Titan X GPU for 30 minutes–30 hours depending on the task.

**Table 2:** Performance of the best-performing models on the validation set, with standard deviations reported over three replicates. AUROC is area under the receiver operating characteristic curve and AUPRC is area under the precisionrecall curve.

| Task | Metric | Performance |
| --- | --- | --- |
| OPENBOND | accuracy | 0.9707 ± 0.0002 |
| | AUROC | 0.9883 ± 0.0001 |
| | AUPRC | 0.9391 ± 0.0006 |
| FRAGMENT | accuracy | 0.9238 ± 0.0012 |
| | AUROC | 0.9652 ± 0.0010 |
| | AUPRC | 0.8496 ± 0.0049 |

## 4 Experiments

### 4.1 Performance of Models in Selecting Actions

Both the OPENBOND model and FRAGMENT model had high performance on validation sets compared to a random baseline, with AUPRC being 0.94 and 0.85 respectively in being able to discriminate between correct (or near-native) and decoy actions (Table 2). To more rigorously access the accuracy of the FRAGMENT evaluation model, we measured how frequently the model scores could be used to select a near-native action out of all potential sampled options (Figure 5). This is a much larger set of options than the select decoys used during training, typically with hundred of choices. The model selected the near-native option 13.1% of the time, compared to 2.6% of the time from a random baseline. Near-native corresponds to nearly exactly matching the action taken within a known ligand; this metric is challenging because in many cases, multiple actions may still generate acceptable ligands so perfect performance is likely impossible and undesirable.

### 4.2 Quantifying the physiochemical properties of generated molecules

Gaining the trust of medicinal chemists has been a challenge for generative models, as molecules produced can be difficult to make in the lab and do not always conform to drug-like physiochemical properties. These aims encompass many separate properties, so we selected 12 metrics both traditionally relevant to medicinal chemists (polar surface area, logP, synthetic accessibility) and some of our own (docking score, number of unsatisfied hydrogen bond donors, torsion energy).

We generated ligands for 85 different proteins that were in the test set, using fragments from real ligands (PDBBind) as starting points. We then measured the predicted physiochemical properties of these ligands across the 12 different metrics. We compared these ligands to the corresponding complete ligands from the PDBBind database. We also generated ligands using a state-of-the-art physics-based scoring functions (Glide) to select actions in place of our models (Friesner et al., 2004). These physics-based scoring functions have been applied previously for ligand optimization tasks, and thus we chose to use this as a baseline of performance. We used the molecular weight of the ligand in the protein-ligand complex from the PDB as the molecular weight goal to determine when to stop adding fragments.

Molecules generated using our models closely matched ligands in the dataset of known ligands across many key properties (Figure 3, Figure 4). These include ligand-only properties, such as logP and synthetic accessibility, as well as properties that take into account the 3D conformation in the binding pocket such as number of unsatisfied hydrogen bond donors and calculated torsion angle energy. One exception is that our ligands had lower absolute formal charge, which may indicate lack of ionic interactions with the protein pocket relative to known ligands. This will be investigated further in future work.

**Figure 3:**
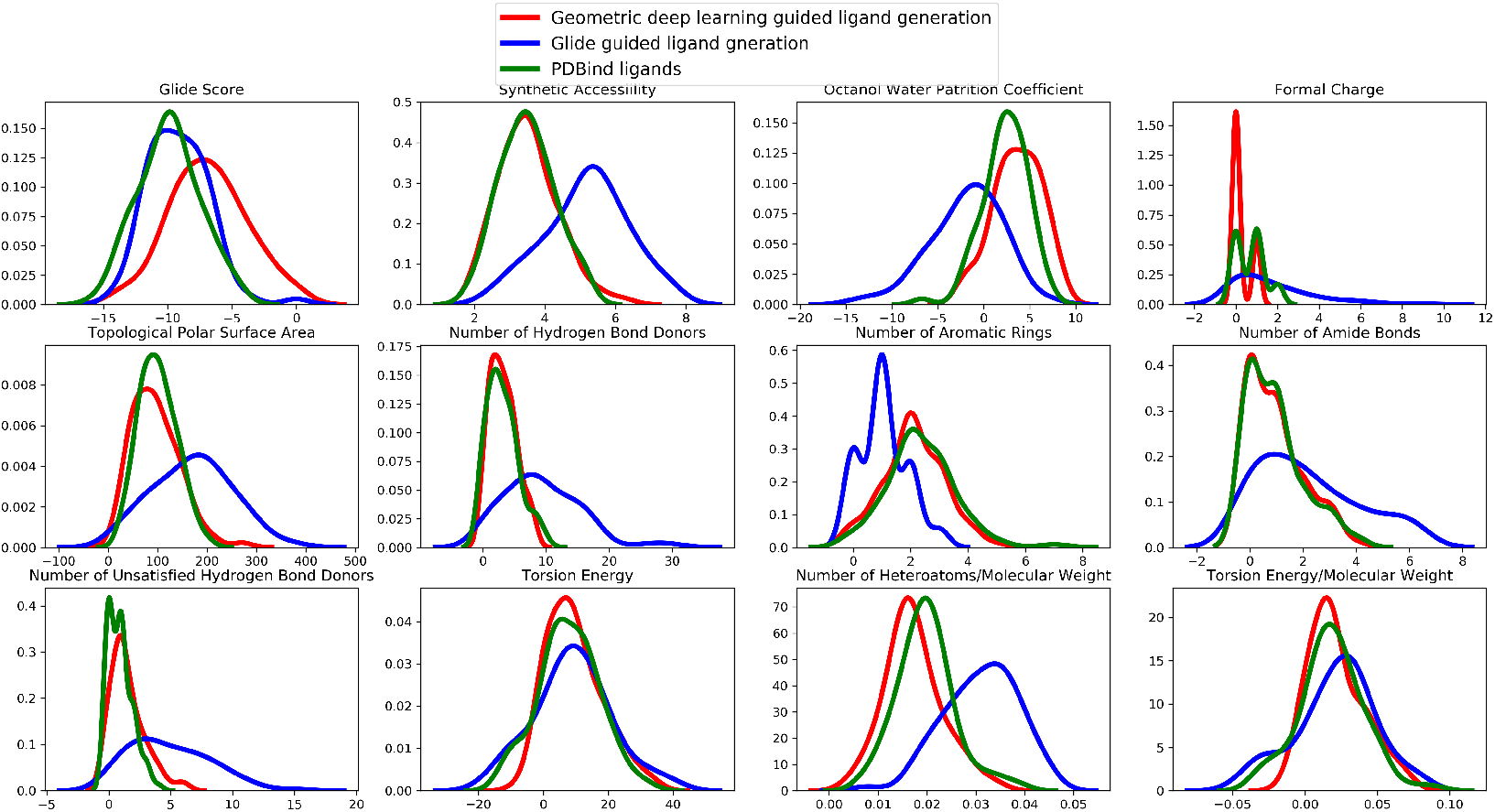
Generated ligands were similar to real ligands across many physiochemical properties. 85 ligands using test set protein pockets were generated using our method. The property distributions of the generated ligands (red) were compared against ligands generated with a physics-based scoring function called Glide (blue) and known ligands in the PDBBind dataset (green). The molecular weight of reference ligand in the PDBBind dataset was used as the molecular weight cutoff to determine when to stop adding fragments. Ligands were scored based upon the Glide score is the docking score after minimization. Torsion energy is computed using Schrodinger structure tools. Other properties computed by RDkit. Formal charge is the absolute value of the formal charge.

**Figure 4:**
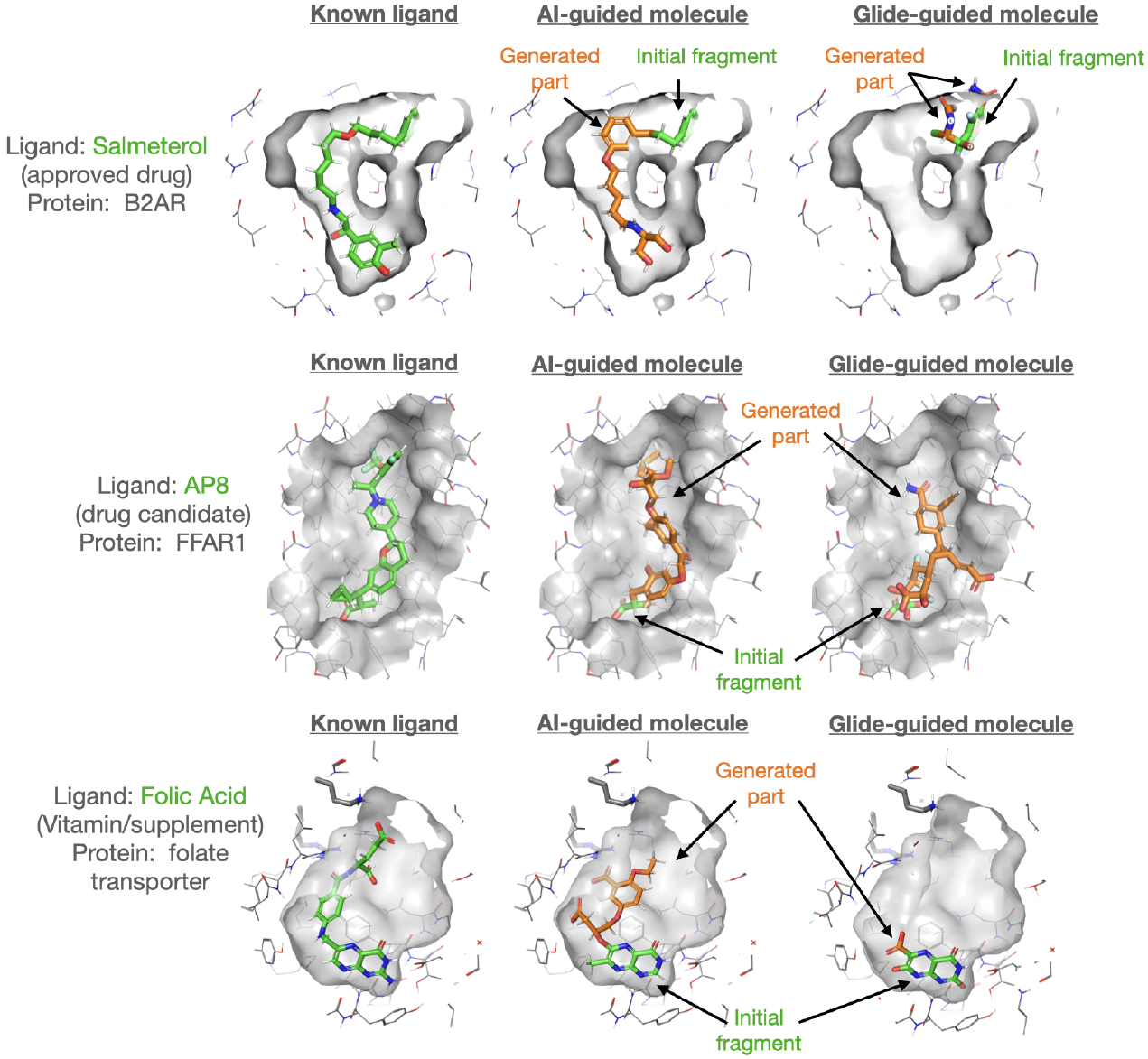
Comparison of ligands generated using different methods. We generated ligands by greedily selecting the top scoring actions until the molecular weight of the ligand hit a target. We compared ligands (left) from that were known binders from the PDB, (middle) ligands generated by our trained models, and (right) ligands generated by selecting actions with a physics-based scoring function called Glide. As seen in the top row and bottom row, for the physics-based scoring function the expansion process occasionally terminated before the goal molecule weight was reached, as there were no valid locations remaining to add to.

In contrast, molecules generated using the state-of-the-art physics-based scoring functions had significant differences from known ligands in our dataset, being more highly charged and more polar overall (Figure 3). Additionally, these ligands contained a higher number of unsatisfied hydrogen bond donors, an energetically unfavorable property frequently used to flag unpromising ligands in virtual screening projects (McDonald & Thornton, 1994; Fischer et al., 2021). However, these molecules did tend to have high docking scores, which is unsurprising given that this property is assessed by the same scoring function used to greedily select actions. This highlights the discrepancy between physics-based docking scores and predicted affinity, although the two are sometimes conflated. How tightly a molecule binds depends on how strongly it can interact with the protein pocket relative to strength of interaction with water (solvation). However, many physics-based scoring functions do not account for solvation effects as it is computationally challenging. Our agent is able to learn this information implicitly because it is trained on a dataset comprised of known binders, molecules that interact with the protein pocket more strongly than with water.

What about exploration of the pocket space? Growing a ligand by sequential addition of fragments benefits from planning ahead capability of our trained models, particularly to avoid running into dead-ends. By qualitative observation, molecules generated using our trained models tended to fill out the pocket and adopt a similar shape as known ligands (Figure 5). With the physics-based scoring function (which has as no planning capability), the expansion process occasionally terminated before the goal molecule weight was reached, as there were no hydrogen atoms remaining to add to (Figure 4).

### 4.3 Interpretability

We next tested how well our models captured basic physics and chemical intuition, factors that allow generalization to unseen protein binding sites. First, using our trained fragment-scoring model, we computed scores for a set of ligand-pocket states that varied only in the dihedral angle of an attached methyl fragment (Figure 6). The overall pattern of scores looked remarkably similar to a dihedral potential energy diagram, with the best score corresponding the most energetically favorable conformation. Next, we tested whether the model learned to account for interactions between the ligand and protein pocket. For example, we observed that the model would successfully place a charged carboxylate fragment to form ionic interactions with an arginine sidechain (Figure 7). When the arginine was mutated to a non-polar leucine, the model would instead place non-polar fragments demonstrating a basic awareness of favorable ligand interactions. Finally, by not explicitly defining fragment types in our model, but rather learning underlying patterns of interactions and geometries, we can introduce new fragment types without retraining (Figure 8). These unseen fragments still received reasonable scores.

## 5 Conclusions

The drug discovery process is long, expensive, and unpredictable. The time it takes to develop a new drug ready to be administered in humans can take 12-15 years (Liszewski, 2006). A major task in drug discovery is searching the chemical space for the molecules that best optimize a set of complex properties like binding affinity, solubility, and synthesizability. In this work, we present a method that can efficiently expand a small fragment bound to a protein pocket into a larger molecule in this drug-like region of chemical space.

Our method relies on several key ideas. First, we break down the generative process into a sequence of individual actions guided by trained models. This gives our method flexibility in being applied to diverse optimization tasks, whether they require adding a few final functional groups or building new scaffolds. Although the molecule generation is a complex, multi-step process, we train models efficiently using supervised learning. Second, we represent states as point clouds in 3D space, which naturally captures the ligand within the context of a protein pocket. With advances in protein structure prediction, the number of available protein structure models has grown exponentially in the past few years (Jumper et al., 2021). Methods that use this information will also grow more accurate as more protein-ligand structures become available. Third, we train models as scoring functions rather than typical policies. By not encoding our fragment library within the model itself, we can easily vary the fragments for each application and the model can potentially generalize to new unseen fragments without retraining.

However, these results come with several caveats. First, the test protein structures we started with were each determined with a “good” ligand bound (and thus carry information about that solution); in real applications the binding site may not be pre-organized to fit a “good” ligand. This could be ameliorated by introducing flexibility and noise into the protein structures during the training process. Additionally, the diversity of the scaffolds that our method can currently produce is somewhat limited by the diversity of our fragment library, especially in regard to different ring systems. Future work will focus on expanding the fragment library and producing many diverse ligand ideas for each pocket.

In summary, our work is a step toward practical AI tools for medicinal chemists that will accelerate drug discovery and advances in human health.

## Author Contributions

Thanks to Namrata Anand for helpful discussions.

## Acknowledgments

PS was supported by a Graduate Research Fellowship from the US National Science Foundation (NSF).

## A Details on the Architecture

The embedding unit consists of two layers of sequential application of self-interaction, point-convolution, self-interaction (Schütt et al. (2017)), nonlinearity, and point normalization (Eismann et al. (2020b)). For the E(3) equivariant layers of the embedding unit, we restrict the maximum filter rotation order to *l* = 2. At each point convolution, it updates the features associated to a given point *p* based on the features of 50 closest neighboring points in the euclidian 3D space, weighted by their distances to *p*. We express these weighted distances in terms of Gaussian radial basis function (RBF) kernel, as a trainable network of two dense layers with hidden layer of size 12. The number of basis and maximum radius of the Gaussian RBF kernel determines the spatial resolution of the kernel, which we chose to be 12.0Å and 12 respectively for our architecture.

Starting from one hot encoding of the basic element type and ligand/fragment flags as feature channels at input, the first layer of the embedding unit mixes those features and outputs 24 feature channels per rotation order (*l* = 0, *l* = 1, and *l* = 2). The second layer of the embedding unit further mixes those features to output 12 feature channels per rotation order. The 0-th rotation order outputs of this layer are then averaged across all points to be passed as input to the aggregator unit.

The aggregator unit consists of 2 fully connected (FC) layers (followed by ELU activation function Clevert et al. (2016) except for the final FC layer) with hidden dimension of 256.

## B Validation of accuracy of fragment adder

**Figure 5:**
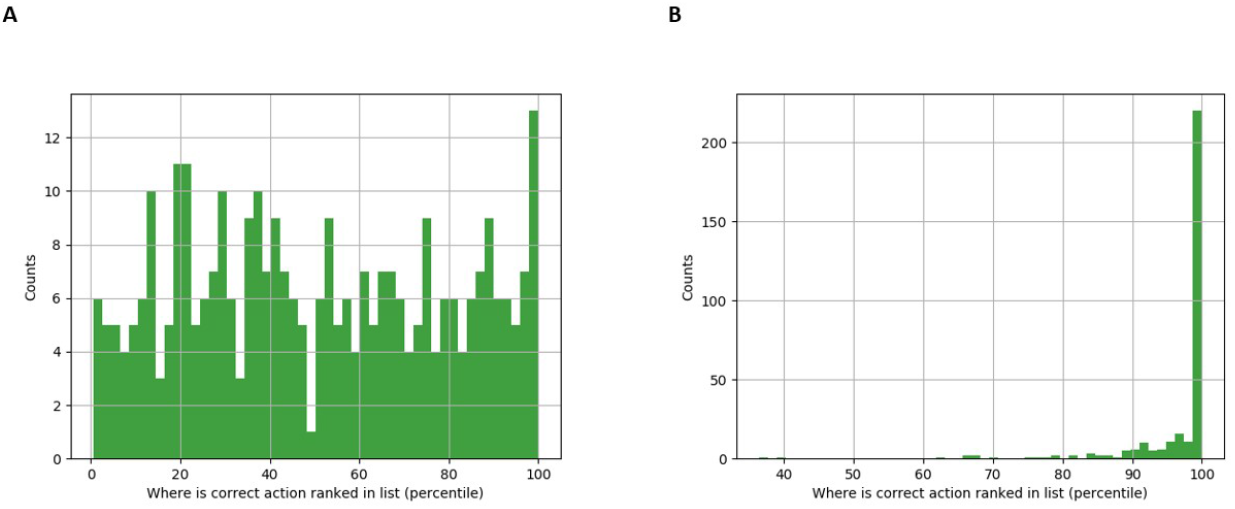
Validation of the accuracy of the Fragment adder: The accuracy of the trained model was evaluated by accessing how frequently it was able to score the “correct” fragment at the correct dihedral angle out of all of the possible intermediates that were generated, relative to randomly selection the correct fragment. Fragmented intermediates from the test set were used to minimize the possibility of data leakage. The trained scoring function was able to substantially out perform random assignment, where the top scoring ligand was ranked in the 73.8th percentile versus 48.0th percentile, given that the software chose to add a fragment to the correct bond. Furthermore, the scoring function was able to correctly identify the “correct” answer 13.1 percent of the time, as opposed to 2.6 from a random selection.

## C Supplementary Figures

**Figure 6:**
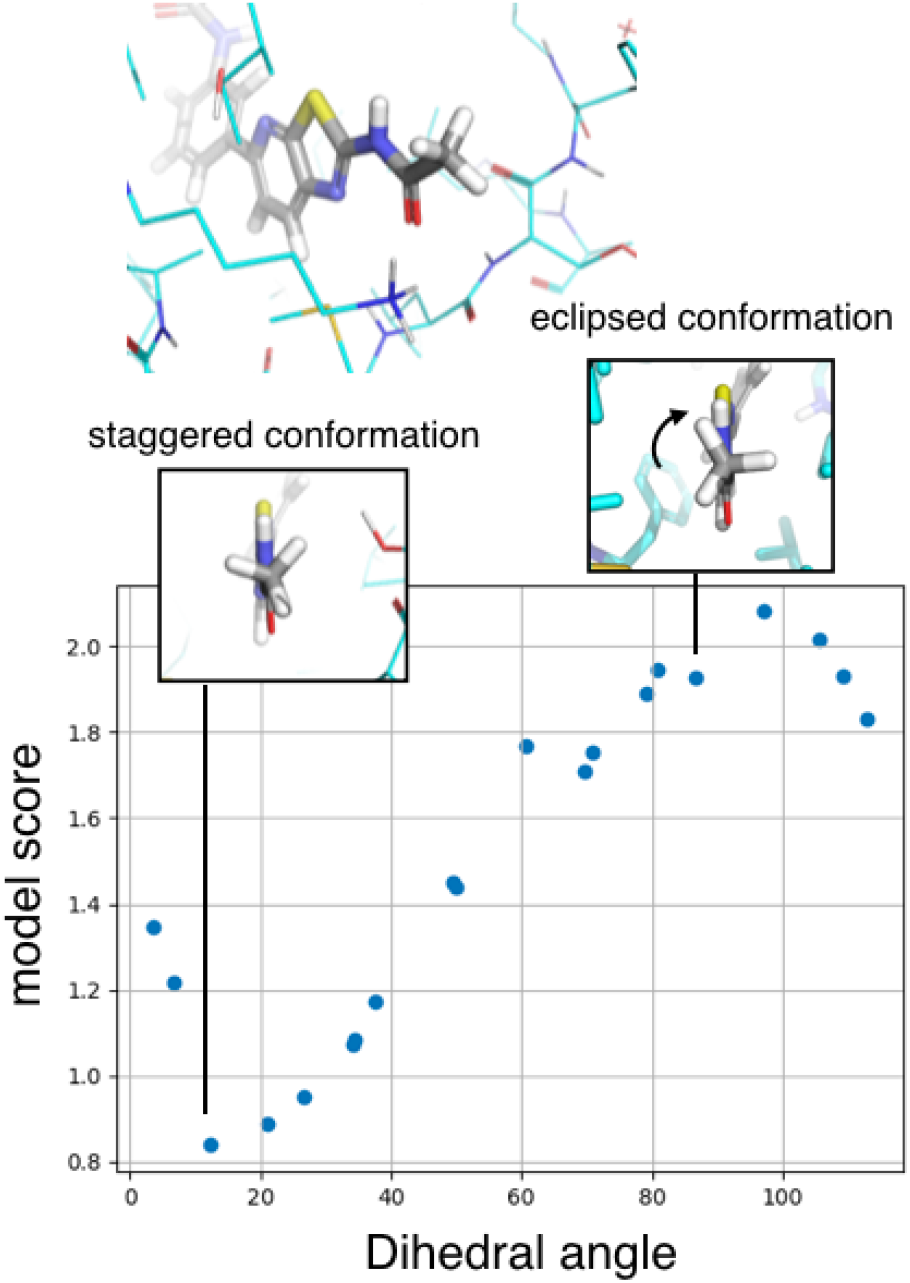
Model learns to approximate physics of chemical geometry. Using our trained fragment-scoring model, we computed score for a set of ligand-pocket structures that varied only in the dihedral angle of the attached methyl fragment. The model predicted the lowest scores (most favorable) corresponding to the staggered conformation, which is known to be the most energetically favorable. The highest scores corresponded to the eclipsed conformation which is the least energetically favorable.

**Figure 7:**
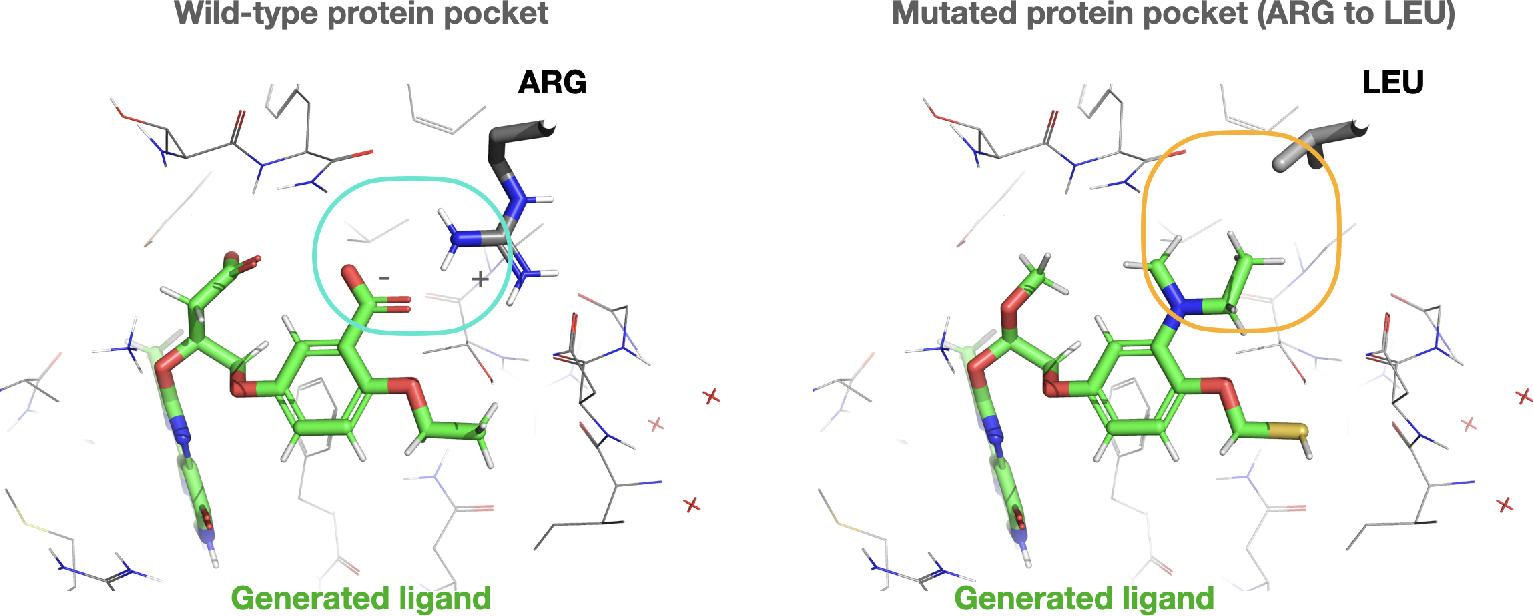
Models learns physics of protein-ligand interactions, and incorporates structure of the protein pocket into selection of chemical fragments. At the folate transporter, the model chose a carboxylate fragment to form an ionic interaction with arginine sidechain (left, blue box). To test this further, we mutated this pocket, by switching the positively charged arginine to a non-polar leucine (right image). When used to generate a ligand for this more non-polar pocket, the model selected more non-polar fragments such as methyl groups to pack favorably against this leucine (orange circle).

**Figure 8:**
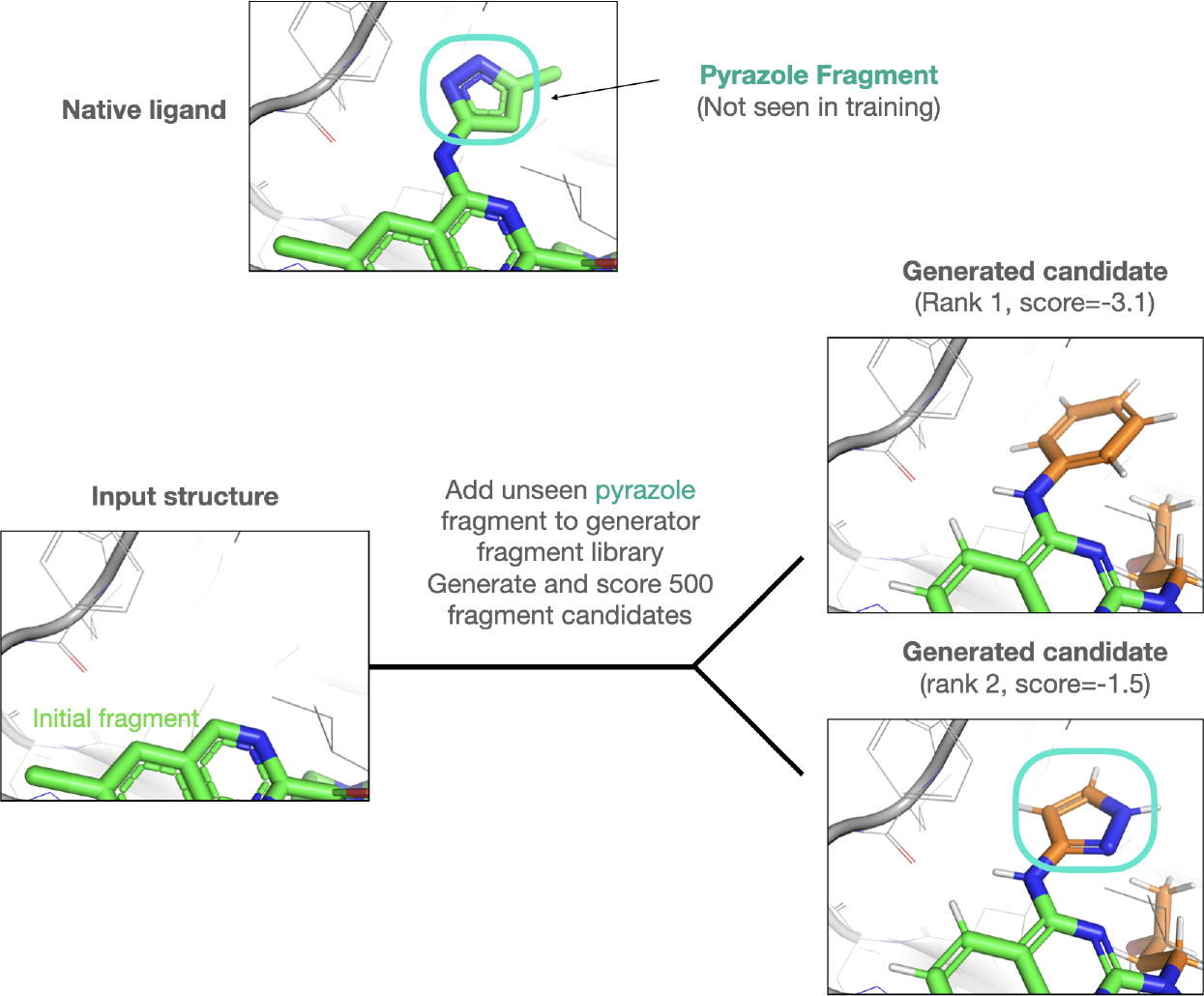
Models can generalize to score fragments not seen during training.

## D Supplementary Tables

**Figure 9:**
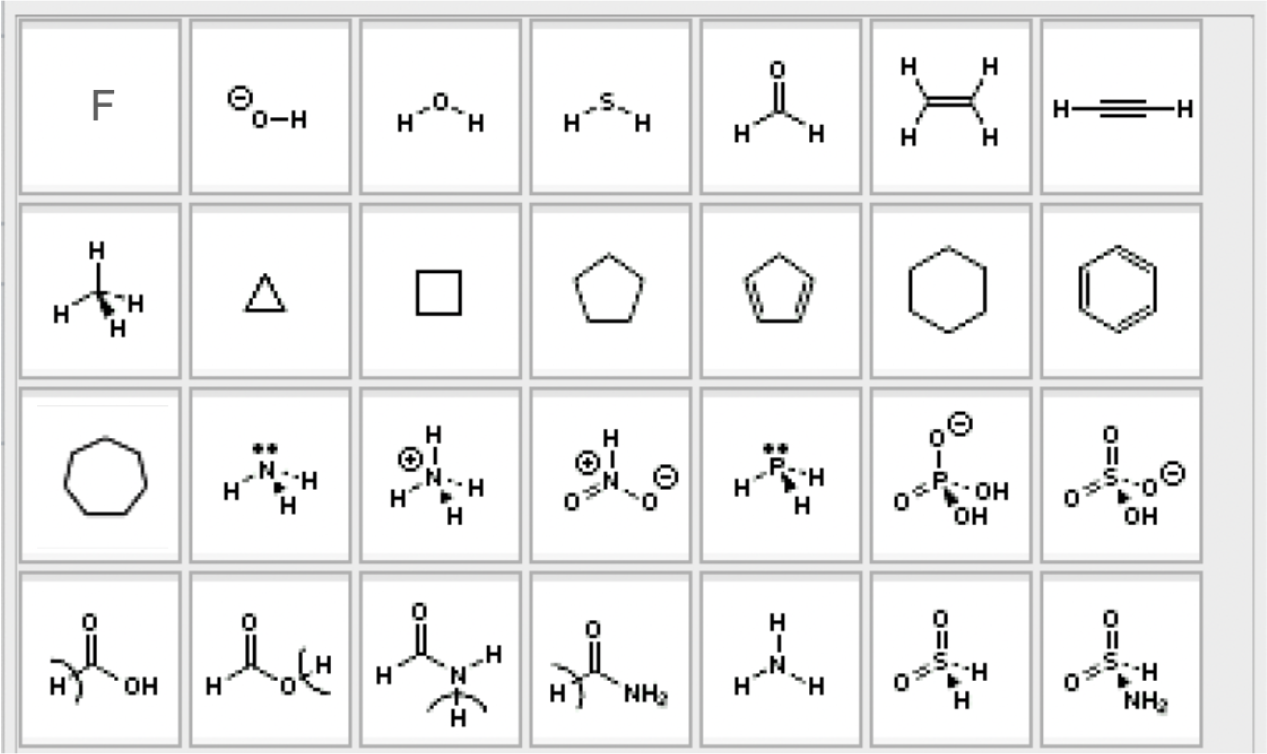
Library of molecular fragments used during training and application.

